## Supplementary Information for "A multi-agent system for spine MRI report generation from multi-sequence imaging"

#### Supplementary Prompts

**System Prompt**

```
system_prompt = """<|begin_of_text|><|start_header_id|system|end_header_id|>.
You are Spine-Agent, a radiology assistant focused strictly on Spine MRI.

Scope and inputs:
- Only discuss Spine MRI and directly related medical context.
- You may be given T1-weighted, T2-weighted, or both.
- Use auxiliary data as supportive context; always validate against the images
    .

Primary tasks:

1) Report Generation
- Produce a structured radiology report with professional tone.
- Summarize key actionable findings in the Impression.

Style and safety:
- Keep responses clear, concise, and focused on the provided Spine MRI.
- Do not fabricate unseen sequences, planes, or prior comparisons.
- Add an appropriate medical caution when relevant."""
```

**Report Generation Prompt**

```
agent_report_generation_prompt = """<image>(1)
If provided, you may also receive auxiliary data delimited by special tokens:
- Disease condition prediction results inside <diagnosis> and </diagnosis>. 0
    represents absence of a disease type, 1 represents presence of a disease
    type.
- Top-1 similar case report inside <similar_case_report> and </
    similar_case_report>

Use auxiliary data as supportive context only. Always prioritize image
    evidence; if there is a conflict, state the discrepancy clearly. Do not
    copy text verbatim from the similar report

<diagnosis>{{}}</diagnosis> <similar_case_report>{{}}</similar_case_report>
    Please analyze these MRI scans and generate a comprehensive radiology
    report including all findings, measurements, and clinical observations.

The templates below are the standard templates for reporting MRI of the
    cervical, thoracic, and lumbar spine. If intravenous contrast was
    administered, enhancement is usually described in the subcomponents of the
     template.

[LUMBAR SPINE TEMPLATE]
FINDINGS:
ALIGNMENT: Normal - Description of the alignment of vertebra in the l-spine.
```

```
MARROW: Normal - Description of the bone marrow and any lesions in the
    vertebra.
DISCS: Discs are normal in height and signal intensity. - Description of the
    inter-vertebral discs, particularly their height loss and desiccation or
    abnormal signal.
CORD: Conus ends normally at L1-L2. Visualized cord and cauda equina are
    normal. - Description of the conus and cauda equina including
    intramedullary and intradural extramedullary lesions.
PARAVERTEBRAL SOFT TISSUES: Normal - Description of findings that are outside
    of the vertebra and spinal canal in the visualized organs and soft tissues
    .
AXIAL DISCS, DURAL COMPRESSION & FORAMINA: - The individual anatomic levels
    below are used to describe primarily the spinal canal, lateral recess, and
     neural foramen at each anatomic level and the degree of compromise as
    well as the etiology for that compression/narrowing.
L1-2: No central or foraminal stenosis. Facets are normal.
L2-3: No central or foraminal stenosis. Facets are normal.
L3-4: No central or foraminal stenosis. Facets are normal.
L4-5: No central or foraminal stenosis. Facets are normal.
L5-S1: No central or foraminal stenosis. Facets are normal.
IMPRESSION: - The impression is a concise restating of the clinically
    important findings of the report as well as the interpretation of those
    findings which may include
specific diagnoses. The impression is also used to describe important items
    that are not present but that need to be understood by the treating
    provider to determine the next course in clinical management.
Provide these diagnoses or issues as an enumerated list.
1.
2.

[THORACIC SPINE TEMPLATE]
FINDINGS:
ALIGNMENT: Normal - Description of the alignment of vertebra in the l-spine.
MARROW: Normal - Description of the bone marrow and any lesions in the
    vertebra.
DISCS: Discs are normal in height and signal intensity. - Description of the
    inter-vertebral discs, particularly their height loss and desiccation or
    abnormal signal.
CORD: Visualized spinal cord is normal in signal and size. - Description of
    the conus and cauda equina including intramedullary and intradural
    extramedullary lesions.
PARAVERTEBRAL SOFT TISSUES: Normal - Description of findings that are outside
    of the vertebra and spinal canal in the visualized organs and soft tissues
    .
SPECIFIC LEVELS: Abnormal levels are described separately below. - Since
    pathology in the thoracic spine is less common the individual anatomic
    vertebral levels are usually not
enumerated. Use this section to describe the specific levels that are
    pathologic, particularly in terms of: spinal canal, lateral recess, and
    neural foramenal compromise.
ALL OTHER LEVELS: No central or foraminal stenosis. Facets are normal.
IMPRESSION: - The impression is a concise restating of the clinically
    important findings of the report as well as the interpretation of those
    findings which may include
specific diagnoses. The impression is also used to describe important items
```

```
    that are not present but that need to be understood by the treating
    provider to determine the next course in clinical management.
Provide these diagnoses or issues as an enumerated list.
1.
2.

[CERVICAL SPINE TEMPLATE]
FINDINGS:
ALIGNMENT: Normal - Description of the alignment of vertebra in the l-spine.
MARROW: Normal - Description of the bone marrow and any lesions in the
    vertebra.
DISCS: Discs are normal in height and signal intensity. - Description of the
    inter-vertebral discs, particularly their height loss and desiccation or
    abnormal signal.
CORD: Visualized spinal cord is normal in signal and size. - Description of
    the conus and cauda equina including intramedullary and intradural
    extramedullary lesions.
PARAVERTEBRAL SOFT TISSUES: Normal - Description of findings that are outside
    of the vertebra and spinal canal in the visualized organs and soft tissues
    .
AXIAL DISCS, DURAL COMPRESSION & FORAMINA: - The individual anatomic levels
    below are used to describe primarily the spinal canal, lateral recess, and
     neural foramen at each anatomic level and the degree of compromise as
    well as the etiology for that compression/narrowing.
C2-3: No central or foraminal stenosis. Facets are normal.
C3-4: No central or foraminal stenosis. Facets are normal.
C4-5: No central or foraminal stenosis. Facets are normal.
C5-6: No central or foraminal stenosis. Facets are normal.
C6-7: No central or foraminal stenosis. Facets are normal.
C7-T1: No central or foraminal stenosis. Facets are normal.
IMPRESSION: - The impression is a concise restating of the clinically
    important findings of the report as well as the interpretation of those
    findings which may include specific diagnoses. The impression is also used
     to describe important items that are not present but that need to be
    understood by the treating provider to determine the next course in
    clinical management.
Provide these diagnoses or issues as an enumerated list.
1.
2.

Please first identify whether the MRI scans are of the cervical, thoracic, or
    lumbar spine, and then use the corresponding template to generate the
    report. If the classification result suggests a region, still verify on
    the images.
<report_generation>"""
```

**Visual Input Specification** — **Note 1**

The <image>(1) tag represents a multi-modal input payload consisting of:

- **Primary MRI Series:** Full-stack T1/T2/others sequences.
- **Segmentation Result:** High-resolution crops of "key-regions" identified by the Segmentation Agent to ensure focus on pathologic levels.

### Supplementary Figures

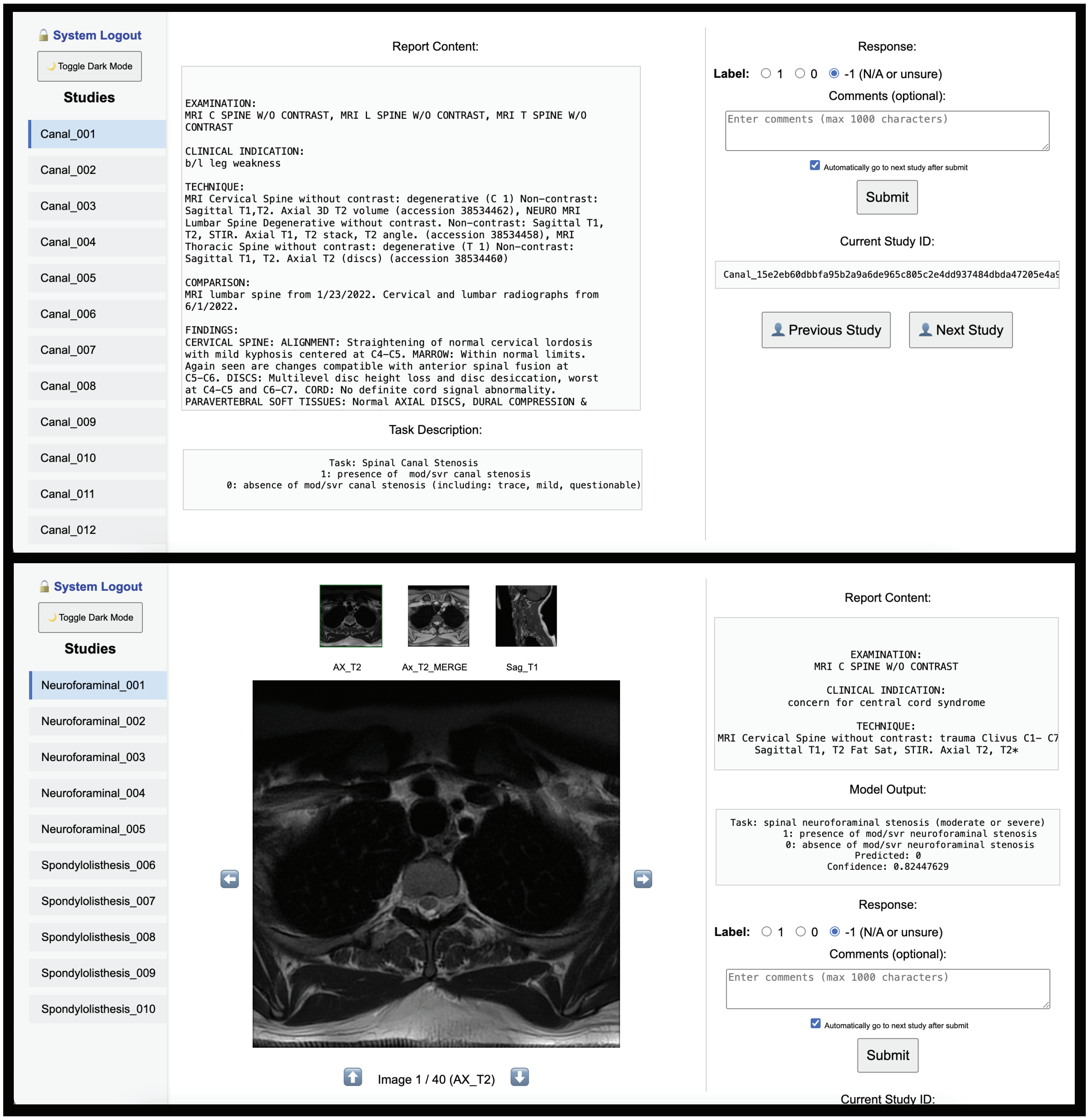


**Supplementary Figure 1: Screenshots of the manual annotation interface** used by radiologists to review and label selected studies from the UW Medical Center dataset. These annotations were used for reliable evaluation of *SpineAgent* and baseline methods.

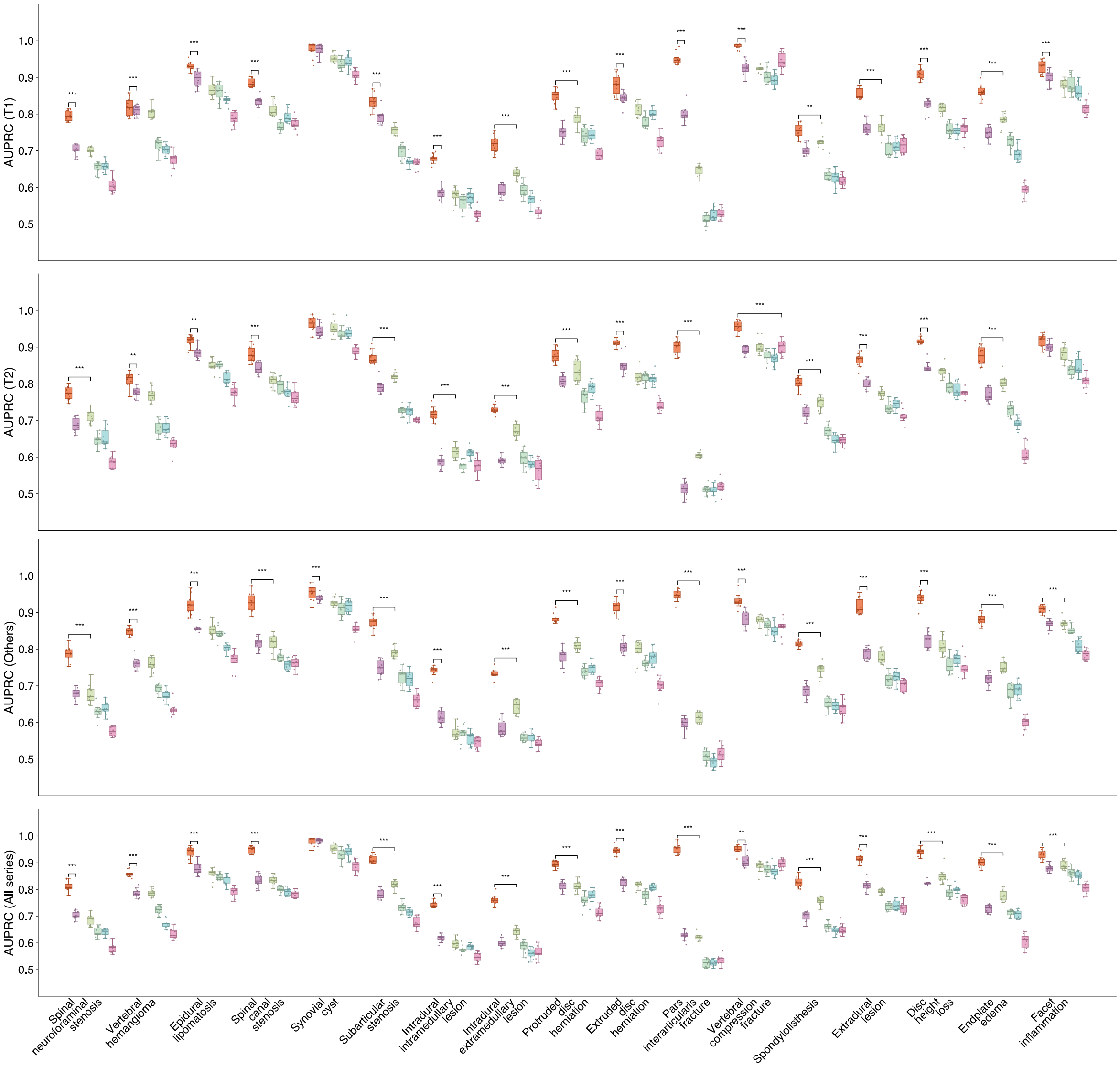


**Supplementary Figure 2: AUPRC comparison** between *SpineAgent* Diagnosis Agents and competing models across different settings, trained on only T1-weighted, only T2-weighted, non-T1/T2 (other) sequences, or all sequences combined. *SpineAgent* consistently achieves superior performance across all settings.

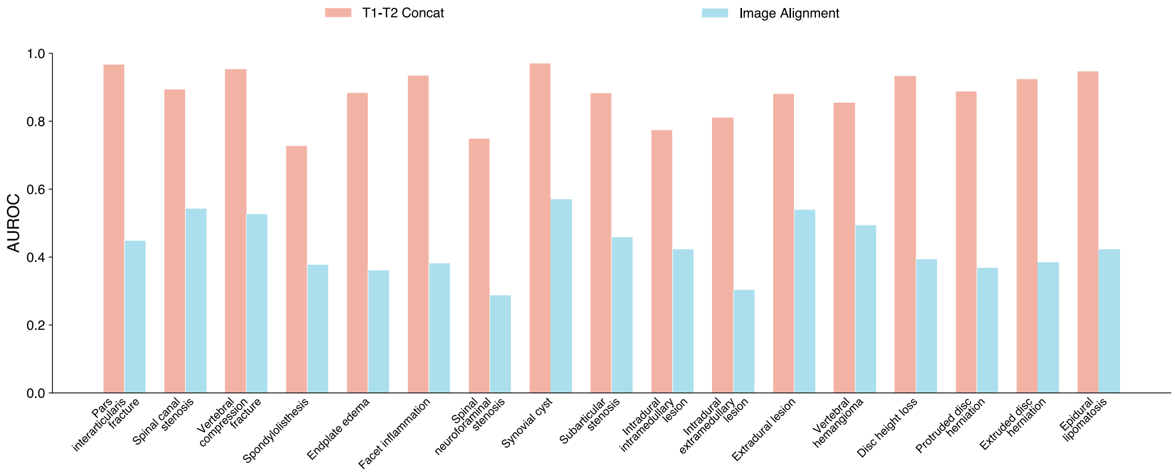


**Supplementary Figure 3: Evidence that image-to-image bi-modal alignment is suboptimal.** Comparison of AUROC scores for combining *SpineFM*'s T1- and T2-weighted representations via direct feature concatenation versus after a CLIP-style alignment between T1- and T2-weighted imaging modalities. Direct image-to-image alignment yields lower performance than simple concatenation, suggesting challenges in aligning heterogeneous MRI sequences.

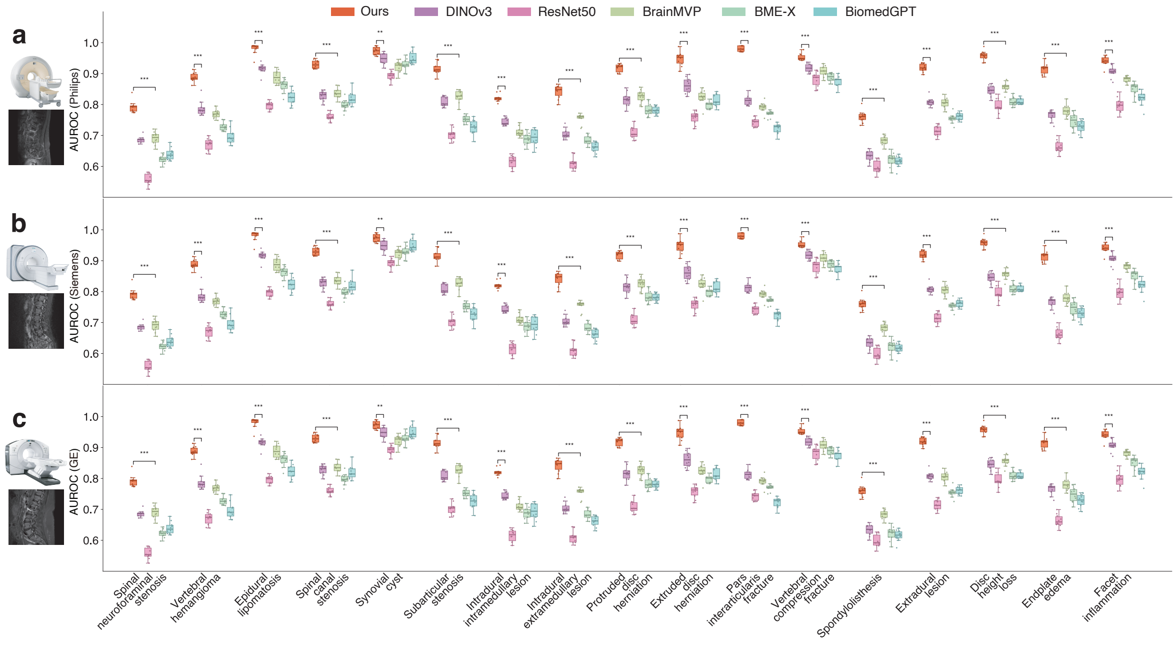


**Supplementary Figure 4: Cross-manufacturer performance** of *SpineAgent* and baseline methods measured by their AUROC scores at the individual condition level. *SpineAgent* consistently outperforms competing methods, demonstrating the robustness and generalization benefits of the pretrained foundation model *SpineFM* across heterogeneous scanner manufacturers.

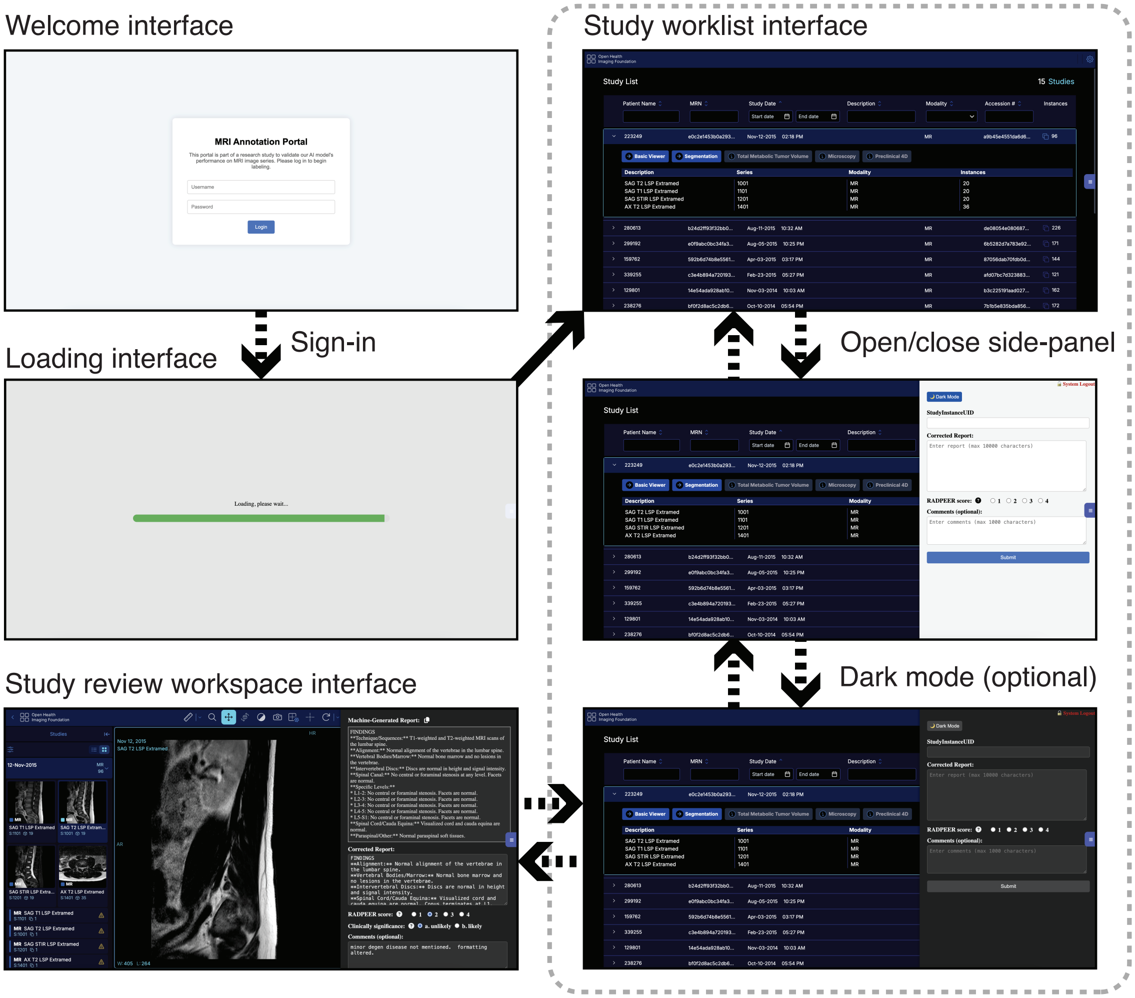


**Supplementary Figure 5: Screenshots of the report-review interface** used by radiologists to evaluate generated reports, provide corrected versions, assign RADPEER scores, and optionally record case-specific comments.

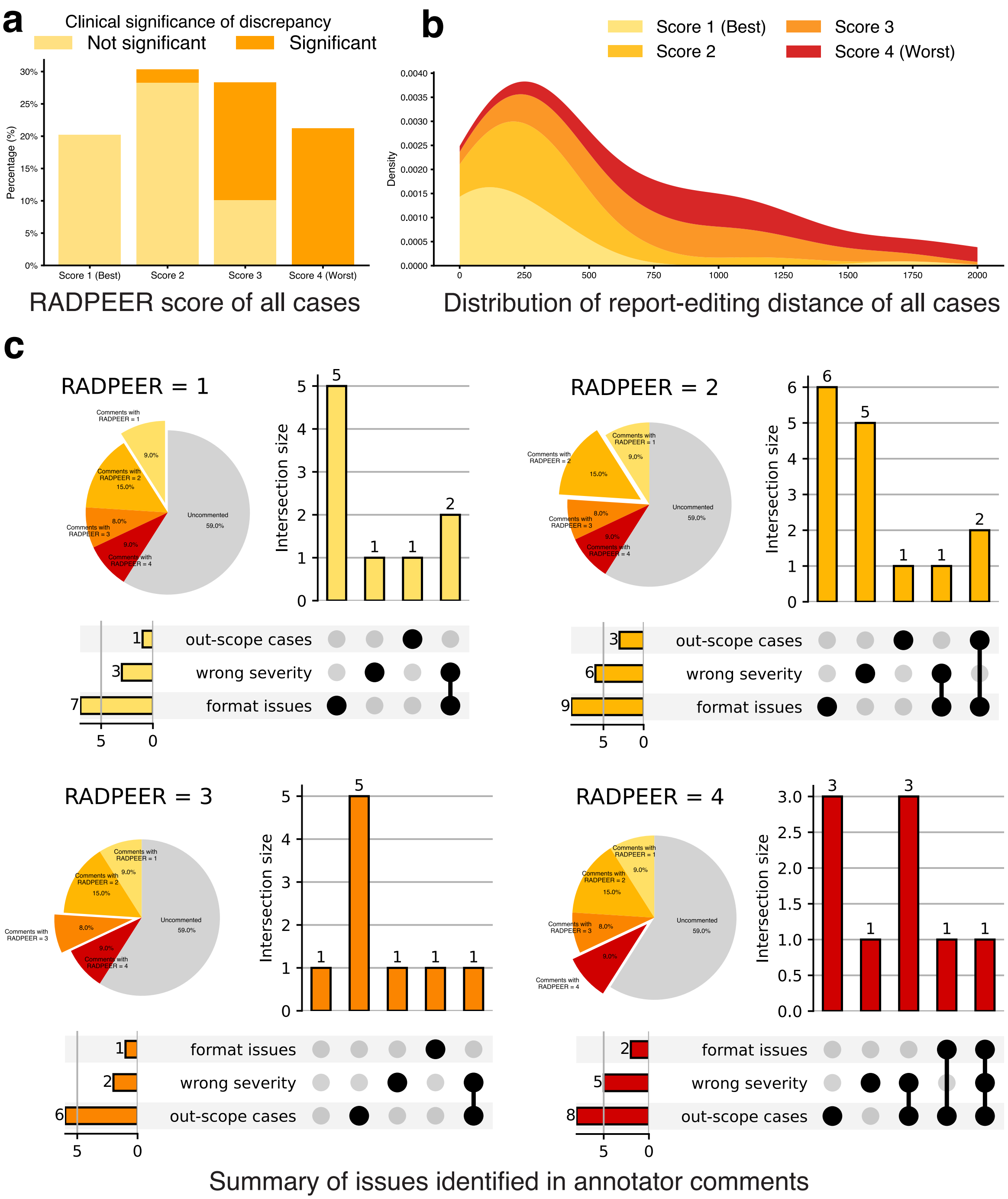


**Supplementary Figure 6: Summary of radiologists feedback across randomly selected** 100 **cases**, including both in-scope and out-of-scope cases. **a,** RADPEER score distribution. **b,** Distribution of editing distances between generated reports and radiologist-corrected versions. **c,** Summary of issues identified in radiologist comments during case review and annotation, reflecting some of the limitations of the current version of *SpineAgent*. Limited label coverage (that is, out-of-scope cases) is the primary contributor to worse RADPEER scores (like 4 or 3).
